# A stretching mechanism evokes mechano-electrical transduction in auditory chordotonal neurons

**DOI:** 10.1101/2025.06.27.662008

**Authors:** Atitheb Chaiyasitdhi, Manuela Nowotny, Marcel Van der Heijden, Benjamin Warren

## Abstract

Insects’ sound and vibration detection including proprioception rely on the scolopidium—a mechanosensory unit enclosing the sensory cilium of chordotonal organ neurons. The cilium, enclosed by a scolopale cell, contains mechanosensitive ion channels with the ciliary tip embedded in a cap. Despite knowledge of the scolopidial structure in multiple insects, the mechanism by which mechanical force elicits transduction remains speculative. We examined scolopidia in the auditory Müller’s organ of the desert locust and present a comprehensive three-dimensional (3D) ultrastructure of a scolopidium using Focused Ion Beam Scanning Electron Microscopy (FIB-SEM). Next, we characterised sound-evoked motions of Müller’s organ and the scolopidium using Optical Coherence Tomography (OCT) and high-speed light microscopy. Finally, we measured transduction currents via patch clamp electrophysiology during mechanical stimulation of individual scolopidia. By combining ultrastructure, sound-evoked motions, and transduction current recordings, our finding suggests that the scolopidium is activated best by stretch along the ciliary axis.

## 1. Introduction

All animal ears share the same fundamental purpose of converting sound-evoked vibrations into electrical signals^1^. In insects, this process, known as mechano-electrical transduction (MET), occurs in the distal ciliated endings of chordotonal organ neurons^2^. The cilium is enclosed in an evolutionarily conserved multicellular structure, the scolopidium^3^, where the cilium extends into a receptor lymph cavity made by the scolopale cell^4^. The ciliary tip is embedded in an extracellular cap tightly fused to an attachment cell. This stereotypical structure of a scolopidium is found in all insect chordotonal organs and underlies sound and vibration detection and proprioception^5^. Thus, understanding how the scolopidium functions has widespread implications for insect behaviour.

While it is established that MET takes place within the cilium, the precise mechanism remains speculative^5^. MET channels open in response to mechanical stimulation leading to a depolarisation that triggers downstream spikes. In *Drosophila* chordotonal organs, putative transduction channels, No mechanoreceptor potential C (NompC) and Nanchung-Inactive (Nan-Iav), are localised either side of the ciliary dilation—an enlarged region where the axonemal microtubules are farther apart than the rest of the cilium. Specifically, NompC localises to the distal end of the cilium, while Nan-Iav is found more proximally, extending from the ciliary rootlet to the dilation^6–9^. Early studies proposed two primary models for how these channels open: axial extension (stretching)^10–12^ or lateral compression (pinching) of the ciliary tip^13^. One prominent axial extension model posits that stretching deforms the ciliary dilation, generating shear forces within the ciliary membrane that open the MET channels^10^. In contrast, the lateral compression model is supported by a similar proposed mechanism in hair plate sensilla, where direct force application on the cap evokes a maximal receptor potential^13^. Alongside the two MET channel opening models—axial extension and lateral compression of the ciliary tip— several other working mechanisms have been proposed based on early ultrastructural and electrophysiological studies^5^. Notably, recent experiments on bush cricket’s hearing organ, crista acustica, suggest that the cap may tilt relative to the ciliary rootlet upon sound stimulation^14,15^. Such a tilting motion could activate the MET channels by stretching the ciliary membrane as the cilium bends, by asymmetrical compressing of the ciliary tip, or through a combination of both.

A detailed ultrastructure of the scolopidium is key to understand its function as a mechano-electrical transducer^3,11,16–19^. The first ultrastructural studies of the scolopidium stem from more than 65 years ago^3,19^, and only recently has three-dimensional (3D) reconstruction using Focused Ion Beam-Scanning Electron Microscopy (FIB-SEM), been employed to examine a miniaturised chordotonal organ, the Johnston’s organ of parasitoid wasp *Megaphragma viggianii*^20^. A high-resolution 3D reconstruction that focuses solely on the scolopidium using FIB-SEM could provide new insights into its ultrastructure, especially how each component interacts.

Complementing these structural insights, efforts to understand the sound-evoked scolopidial motion have yielded valuable, albeit indirect, inferences: Stephen and Bennet-Clark’s pioneering stroboscopy of the locust Müller’s organ suggested significant phase lags and anti-phasic motion across the organ, potentially inducing strain in auditory neurons and scolopidia^21^. More recently, Optical Coherence Tomography (OCT) based vibrometry of the bush cricket’s crista acustica indicated complex, multi-modal tilting of the ciliary tip, including 3D elliptical motions^14,15^. Although providing valuable insight, these observations lack the spatial resolution to directly resolve scolopidia. This limitation highlights the critical need for direct visualisation of its sound-evoked motion, and measurement of transduction currents during direct mechanical stimulation of a scolopidium.

To address these gaps, we investigated scolopidia within Müller’s organ of adult desert locusts, *Schistocerca gregaria*, a model organism that uniquely enables recordings of transduction channel currents from auditory neurons in an adult insect^9^. The desert locust’s tympanal ears are located on its first abdominal segment (Fig. 1a-b). The locust’s hearing organ, Müller’s organ, is directly attached to the inside of the tympanic membrane at an inward fold of the tympanum called the elevated process (Fig. 1b-c). Auditory neuron somata locate mostly in the ganglion, while their dendrites extend to distinct regions of the organ, forming three groups also distinguished by characteristic frequency ranges^9,22^ (Fig. 1c, Extended Data Fig. 1a-c). This study examines scolopidia from Group-III auditory neurons (Fig. 1c-d), which constitute most auditory neurons (∼48 out of ∼80 neurons) in Müller’s organ^23^ and are tuned to 2.5-4 kHz^1^, the frequency ranges associated with wing-beat noise^24^.

**FIGURE 1.**
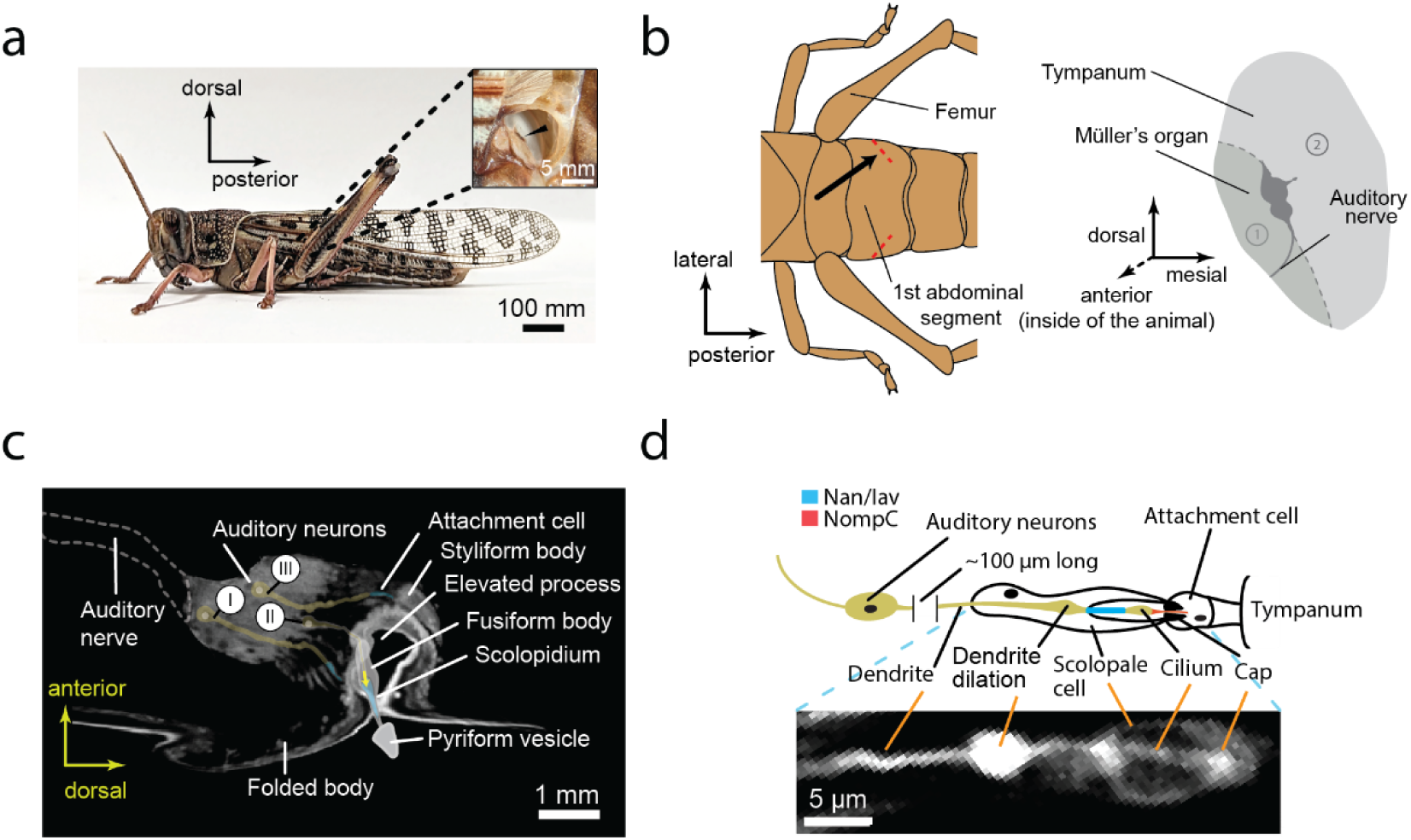
The locust’s ear, Müller’s organ, and the scolodium. **(a)** Locust’s ears are located on the first abdominal segment, hidden behind the wings. Each ear consists of a tympanic membrane with a hearing organ, Müller’s organ, directly attached. External view of the tympanic membrane (inset) shows a dark ridge (arrowhead) where Müller’s organ is located. The anatomical references used in this manuscript are defined based on the axes of the locust’s body unless stated otherwise. **(b)** The surface of the tympanic membrane aligns orthogonally to the antero-posterior axis of the locust body at a slight oblique angle (left, red dashed lines). Illustration of the right ear viewed from inside the body (along the orientation of the black arrow on the left panel) shows Müller’s organ located between the ‘thick’ (1) and ‘thin’ (2) regions of the tympanic membrane (right). The auditory nerve projects ventrally toward the ventral nerve cord. **(c)** Schematic of Müller’s organ superimposed on an X-ray micro-CT scan slice shows three groups of auditory neurons. These neurons (yellow) extend their dendrites to distinct regions where the distal ciliated end of the dendrites insert into the attachment cells (blue). The auditory neurons of Müller’s organ are classified based on their maximal frequency sensitivity. Group-I neurons respond to frequencies between 0.5 and 1.5 as well as 3.5 kHz (∼20 cells). Group-II neurons are sensitive to higher frequencies of 10–25 kHz (∼13 cells), while most auditory neurons, belong to Group-III and respond to mid-range frequencies of 2.5–4 kHz (∼46 cells). The yellow arrow marks a typical location of a scolopidium at the insertion of the scolopale cell into the attachment cells. **(d)** Schematic of a scolopidium indicating locations of the mechano-electrical transduction (MET) channel candidates, NompC (red) and Nanchung-Inactive (blue), near the proximal and distal ends of the ciliary dilation (top). The bottom panel shows fluorescent micrograph of the corresponding region of the dendrite and the scolopidium. The auditory neuron is filled with Dylight-488 conjugated streptavidin-neurobiotin (bottom).

Here, we combined FIB-SEM to resolve the 3D ultrastructure of a scolopidium, OCT and high-speed microscopy to examine sound-evoked motion at both the organ and individual scolopidium levels, and direct mechanical stimulation of the scolopale cap, where the ciliary tip is anchored, whilst simultaneously recording transduction currents. Understanding the structural and functional dynamics of scolopidia in insect auditory transduction is not only key to deciphering insect hearing but offers insights into basic principles of ciliary mechanosensation across insects.

## 2. Results

### 2.1. Three-dimensional (3D) ultrastructure of a scolopidium

To reconstruct a 3D ultrastructure of a scolopidium, we performed Focused Ion Beam-Scanning Electron Microscopy (FIB-SEM) at the interface between the scolopale cells and the attachment cells, where the scolopidia are located (Fig. 2a-b, see Supplementary Video 1) (N = 2, n = 3). In this region, the scolopale cell forms a protrusion that encloses the scolopale sheath with approximately half of the sheath inserting deep into the attachment cell. The scolopale sheath, previously described as composed of 5-7 partially fused, actin-rich concentric rods, is contained within the scolopale cell^3,25^. In early ultrastructural studies, the distal region of the scolopidium was thought to end with the extracellular porous cap^3,10,11,17^; however, our 3D reconstruction reveals a thin layer of fibrous filaments, likely actin filaments based on phalloidin staining of a similar structure in *Drosophila* scolopidia^26^, within the attachment cell at the interface between the attachment cell and the scolopale sheath (Fig. 2c, arrowhead). Upon examining series of longitudinal sections along the length of the scolopidium, this fibrous material covers the distal end of the ciliated dendrite and the cap like a cone-shaped ‘lid’ (Fig. 2d, arrowheads). Thus, we describe the ensheathed part of a scolopidium as a cartridge composed of the scolopale sheath and a scolopale lid (Fig. 2l).

**FIGURE 2.**
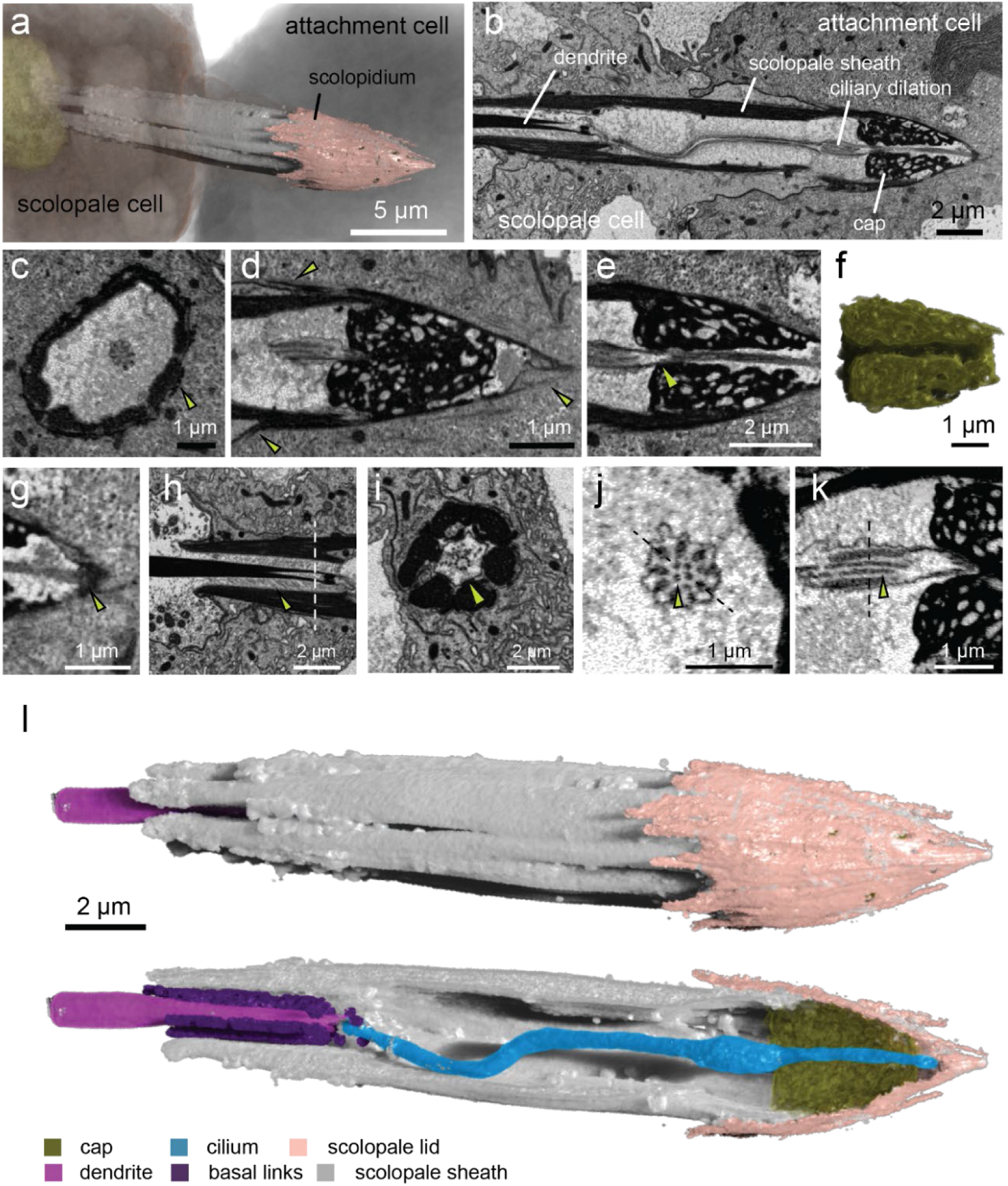
Three-dimensional (3D) ultrastructure of a scolopidium. **(a)** 3D reconstruction at the interface between a scolopale cell and an attachment cell using Focused Ion Beam-Scanning Electron Microscopy (FIB-SEM). **(b)** Longitudinal FIB-SEM slice reveals the dendrite and its ciliated distal end enclosed within the scolopidium sheath; the cilium dilates near its insertion into the porous cap before it terminates in an electron-dense region at the border of the attachment cell. **(c)** Cross section at ciliary dilation shows fibrous filaments inside the membrane of the attachment cell (arrowhead). **(d)** Longitudinal section at the distal region of the scolopale sheath reveals that the fibrous filaments (arrowheads) observed in (c) form a cone-shaped lid covering the cap and the distal region of the ciliated dendrite. **(e)** Longitudinal section at the plane of the cilium shows the narrowing of the central passage, where the cilium enters (arrowhead). **(f)** 3D reconstruction of the cap material reveals a funnel-shaped central passage. **(g)** Termination of the cilium at the extreme end of the scolopale cartridge. **(h)** Longitudinal section at the scolopidium base reveals numerous oblique basal links between scolopale rods and the dendrite. **(i)** Cross section along the dashed line in (h) shows the basal links radiating from desmosome-like junctions. **(j)** Cross section at the ciliary dilation reveals four central fibrils (arrowheads) inside the 9-doublets axonemal structure. **(k)** Longitudinal section along the dashed line in (j) reveals two of the central fibrils along the length of the ciliary dilation. **(l)** 3D model of a complete scolopidium. The auditory neuron in the 3D model is omitted to reveal the oblique basal links.

The scolopale cartridge encloses a tightly sealed receptor lymph-filled scolopale space that contains the ciliated dendrite and the cap (Fig. 2b). Our 3D reconstruction reveal that the cap resembles a truncated cone, with a central passage for the cilium that is funnel-shaped and is distinctly narrow at its entrance (Fig. 2e-f). An early ultrastructural study of Müller’s organ reported that the cilium terminates within the cap^3^; however, we find that the cilium instead extends through the central passage and terminates at an electron-dense region at the tip of the cartridge, in the three scolopidia we imaged (Fig. 2g, Extended Data Fig. 1d). Interestingly, one of our datasets shows a break of the cilium near its terminus, which could have led to the assumption that the cilium terminates in the porous cap (Extended Data Fig. 1e). The cilium’s full extension to its insertion at tip of the cartridge is a prominent, previously undescribed feature with clear mechanical implications.

At the proximal region of the ciliated dendrite, the cilium is anchored through rootlets. Here, numerous basal links connect the dendritic membrane to the base of the scolopale sheath and form an oblique angle of about 50° to the dendrite (Fig. 2h). The basal links show no apparent spacing and are found from the base of the scolopale sheath up to the proximal basal body, where the cilium inserts into the scolopale space. In a cross section orthogonal to the ciliary axis, the basal links appear to radiate towards the dendrite from desmosome-like junctions at the scolopale cell membrane opposing each scolopale rod (Fig. 2i). The basal links and their connections to the base of each scolopale rod have been previously shown^10,11,27^, but not the oblique nature of the links.

Bending occurs consistently at a particular region within the proximal cilium in all our datasets, a feature previously observed in other chordotonal organs^5,12,28^; however, we (and other studies) cannot rule out fixation artifacts being the cause of the bending due to preparation for electron microscopy. In contrast, the distal cilium remains straight (Fig. 2b, l). In the ciliary dilation, where the 9×2+0 axonemal microtubules are farther apart, we found 3-4 central fibrils within the restricted length of the dilation but nowhere else (Fig. 2j-k, Extended Data Fig. 1f-g). While these central fibrils have been mentioned by early ultrastructural studies^3,10^ and observed in crista acustica of the bush cricket, *Mecopoda elongata*^14^, their precise nature remains unclear.

In summary, our high-resolution 3D ultrastructural analysis has redefined the ensheathed part of the scolopidium as a ‘cartridge’ composed of the scolopale sheath and scolopale lid (Fig. 2l). This analysis also revealed the full extent of the cilium (Fig. 2g), the oblique nature of the basal links (Fig. 2h), and the central fibrils in the dilation (Fig. 2j-k). This refined anatomical understanding is crucial for interpreting how forces are transmitted within this mechanosensory unit. Next, we therefore examined sound-evoked motion at the level of Müller’s organ and individual scolopidia to determine how these intricate structures are mechanically stimulated.

### 2.2. Sound-evoked motion of Müller’s organ is tuned differently across its axes

The tympanic membrane vibrates in response to sound pressure changes in the air^29–31^. This sound-evoked vibration is then transferred to Müller’s organ and mechanically activates the scolopidia inside^22,29^. The resulting motion of the organ is complex and frequency-selective, arising from both the tonotopic traveling wave across the tympanic membrane and the organ’s attachment to mechanically heterogenous regions on the tympanum^21,29,31^. Ultimately, however, only the relative motion within the organ can produce forces that serve as the adequate stimulus for auditory transduction.

To identify the source of this relative motion, we measured the sound-evoked vibrations of Müller’s organ along two primary axes (Fig. 3a): (i) antero-posterior (motion orthogonal to the tympanic surface, Fig. 3a-top panel) and (ii) meso-lateral (motion tangential to the tympanic surface, Fig. 3a-bottom panel). The OCT imaging system generated sectional images across the two planes (Fig. 3b) and resolved most of the organ, apart from the region of Group-I scolopidia above the folded body, possibly due to low reflectivity (Fig. 3b). Notably, we could not resolve the Group-III scolopidia along the ventro-dorsal axis—which runs parallel to the dendrite—as the OCT beam was obstructed by either the cuticle or the elevated process.

**FIGURE 3.**
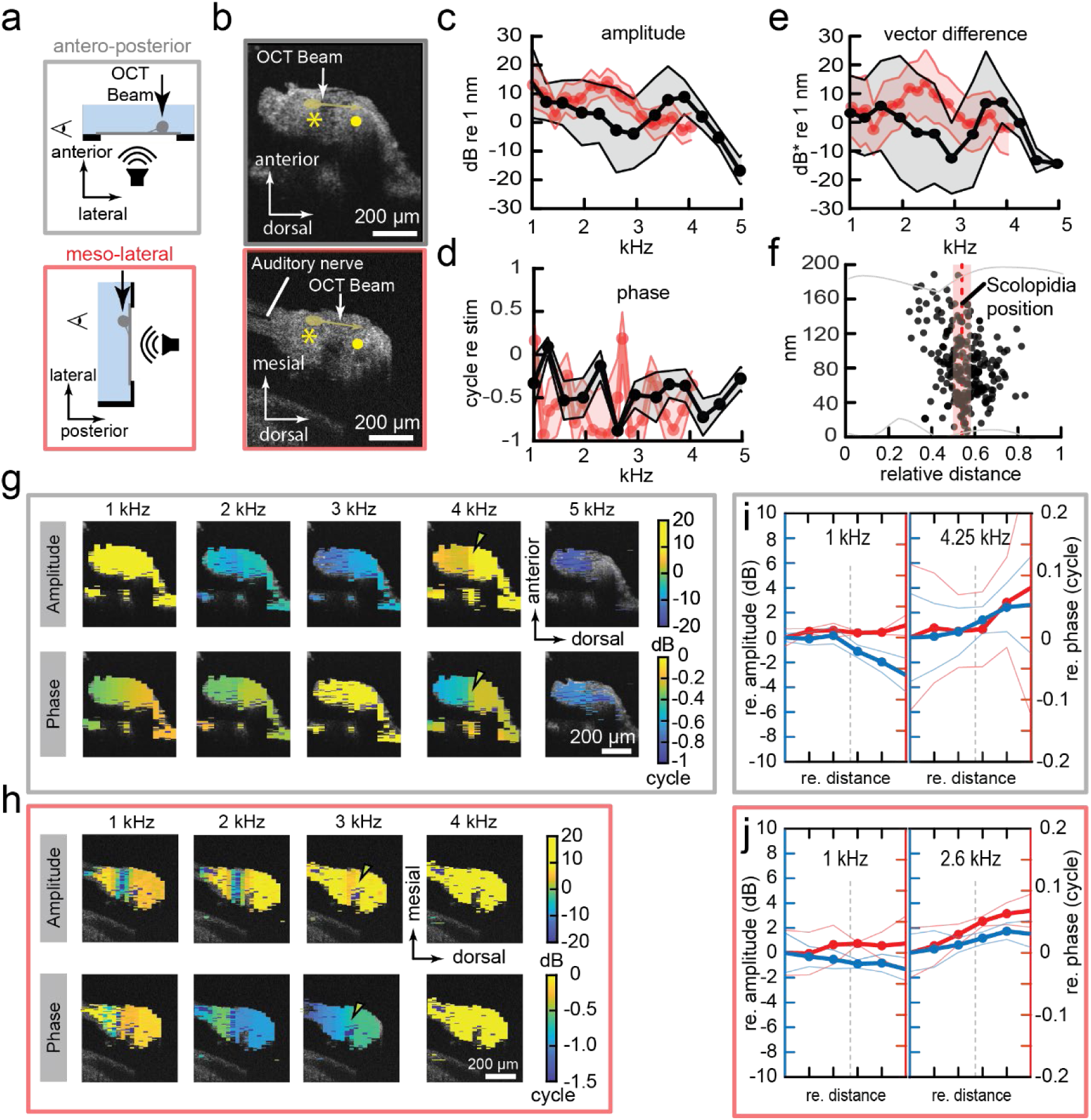
Sound-evoked motion of Müller’s organ. **(a)** Vibrometry measurements were performed using Optical Coherence Tomography (OCT) along the antero-posterior (top) or meso-lateral (bottom) axes of Müller’s organ upon 90 dB SPL sound stimulation, with the external side of the tympanic membrane facing the loudspeaker. Vertical arrows indicate the direction of the OCT beams, and the axes of vibration measured. The eyes represent the observation angles, where 2D OCT sections in (b) were reconstructed. **(b)** OCT sections along the two planes were reconstructed from OCT depth profiles. The symbols * and ● indicate the maximal widths of Müller’s organ at the ganglion and the styliform body, the two important landmarks used in our analysis. Yellow arrows indicate typical orientation of Group-III dendrites. **(c)** Ensemble amplitude and **(d)** phase responses to stimulated frequencies along the antero-posterior (black, N = 3, E = 3) and meso-lateral (red, N = 7, E = 12) axes. **(e)** Vector difference as a function of stimulated frequency, computed by comparing vibration at the position ● relative to *, as shown in (b), along the antero-posterior and meso-lateral axes. **(f)** Normalised positions of Group-III scolopidia between the ganglion and the styliform body (N = 16, E = 16, n = 235). The grey line indicates the outline of Müller’s organ. **(g-h)** Vibration maps as a function of stimulated frequency along (g) the antero-posterior and (h) meso-lateral axes. Arrowheads in (g-h) indicate the area of abrupt transitions in both the phase and amplitudes. **(i-j)** Comparison between relative amplitude (blue) and phase (red) responses from the position * to ● along (i) the antero-posterior (N = 3, E = 3) and (j) meso-lateral (N = 7, E = 12) axes. The relative amplitudes are selected from 1 kHz (minimal relative amplitude) and at the frequencies where the relative amplitude is largest in each axis, 4.25 kHz and 2.6 kHz, respectively. The relative distance is normalised with * as 0 and ● as 1, as shown in (b). The grey dashed line represents the average Group-III scolopidia position obtained from (f). All vibration amplitudes are shown in dB relative to a 1-nm reference vibration. For additional data, see Extended Data Fig. 2 and Source Data Fig. 3.

We first examined the frequency response of Müller’s organ relative to sound stimulation in response to 90-dB-SPL tones (Fig. 3c-d). The vibration was axis-dependent: along the antero-posterior axis, the amplitudes peaked at 3.9 kHz, 9 ± 5 dB re 1 nm (N = 3, E = 3), while the meso-lateral axis showed a distinct peak at 2.4 kHz, 14 ± 5 dB re 1 nm (N = 7, E = 12) (Fig. 3c). There is also relatively large vibration at 1 kHz in both axes, antero-posterior: 14 ± 10 dB re 1 nm (N = 3, E = 3) and meso-lateral: 13 ± 8 dB re 1 nm (N = 7, E = 12). The organ exhibited large phase lag relative to the sound stimulus in both axes (Fig. 3d). Vibration at 1-2 kHz and 3.75 kHz correspond to known resonance modes of the entire tympanum and Müller’s organ region^29^.

To assess the relative motion across the organ, we further computed vector difference, a measure of relative motion that accounts for differences in both vibration amplitude and phase (see Methods), at the styliform body (●, Fig 3b) relative to the ganglion (*, Fig 3b), the region where Group-III scolopidia are located. Quantification of relative motion and vector difference are key to understand where forces that ultimately open MET channels are generated. Vector difference revealed two prominent peaks at 3.9 kHz along the antero-posterior axis, 7 ± 6 dB (N = 3, E = 3), and 2.4 kHz along the meso-lateral axis, 14 ± 11 dB (N = 7, E = 12). The small vector difference at 1.25 kHz in both axes, antero-posterior: 3 ± 11 dB (N = 3, E = 3) and meso-lateral 3 ± 10 dB (N = 7, E = 12), indicates no or very small relative motion in the region of Group-III scolopidia at this frequency (Fig. 3e). Next, we mapped these complex, frequency-dependent vibration patterns onto the locations of Group-III scolopidia.

### 2.3. Relative motion at the site of Group-III scolopidia

We mapped Group-III scolopidia by filling their auditory neurons with the neural tracer neurobiotin, delivered through back-fill of the auditory nerve, then staining with fluorescently-conjugated strepavidin that bound to the neurobiotin^32^. Since ciliated dendrites within the scolopale sheaths were not consistently labelled, we used their dendrite dilations as reliable proxies (Fig. 1d). The position of each dendrite dilation was manually measured relative to the ganglion and the styliform body (Extended Data Fig. 2a-c). These dilations locate at a known distance, 11.1 ± 3.7 µm (N = 20, n = 46), from the ciliary rootlet (Extended Data Fig. 2d). Ensemble normalised positions of the dendrite dilations revealed that Group-III scolopidia are clustered at a relative distance 54 ± 4% (N = 16, E = 16, n = 253) between the ganglion and the styliform body (Fig. 3f).

Having established the location of the Group-III scolopidia, we analysed the 2D vibration profiles to determine if this location corresponds to a region of high relative motion (Fig. 3g-h). Indeed, we observed that while motion was largely homogeneous at low (1 kHz) and high (5 kHz) frequencies, a distinct spatial transition emerged at the intermediate frequencies. Along the antero-posterior axis, an abrupt change in both amplitude and phase appeared from 2 kHz upwards, with the largest transition occurring at ∼4 kHz (Fig. 3g arrowhead, Extended Data Fig. 2e-f). A similar sharp transition was observed in the meso-lateral axis, though its peak occurred at a lower frequency of ∼3 kHz (Fig. 3h, Extended Data Fig. 2g-h). Interestingly, upon examining the vibration profile (Extended Data Fig. 2e, g), the source of vibration appeared to shift with frequency. At lower frequencies (1-2 kHz), the ganglion side of the organ vibrated more strongly (∼2.5-4 dB) than the styliform body, while at higher frequencies the vibration was larger at the styliform body and the tympanum (Extended Data Fig. 2e-h). Crucially, upon superimposing the normalised position of Group-III scolopidia on the frequency response for both axes, the onset of this abrupt transition in amplitude and phase co-localizes precisely with the average position of the Group-III scolopidia (Fig. 3i, j and Extended Data Fig. 2). To understand how this complex relative motion activates the scolopidium, we recorded the sound-evoked motion of individual scolopidia with high-speed light microscopy.

### 2.4. Elliptical motion and stretching of a scolopidium along the ciliary axis

To examine sound-evoked motion at the single-scolopidium level, we captured its high-speed motion during sound stimulation (Supplementary Video 2) and achieved an average acquisition rate of 8,602 ± 289 frames per second (N = 12, n = 48). Upon initial image analysis, we found two prominent motion components (Fig. 4a-b): (i) stretching of the scolopidium (Fig. 4a, t = 0.12 and 0.24 ms) followed by (ii) pulling motion of the scolopale sheath along the ciliary axis towards the attachment cell (Fig. 4a, t = 0.36 ms). The stretching appeared to occur concurrently with a shearing motion. This shear was most apparent with deformation of cell membranes at the cap and the scolopale base (Fig. 4a, yellow lines). To quantify these complex scolopidium motions, we tracked the positions of two clearly visible landmarks: the base of the scolopale sheath, or ‘the scolopale base’ and the extracellular cap (Fig. 4a).

**FIGURE 4.**
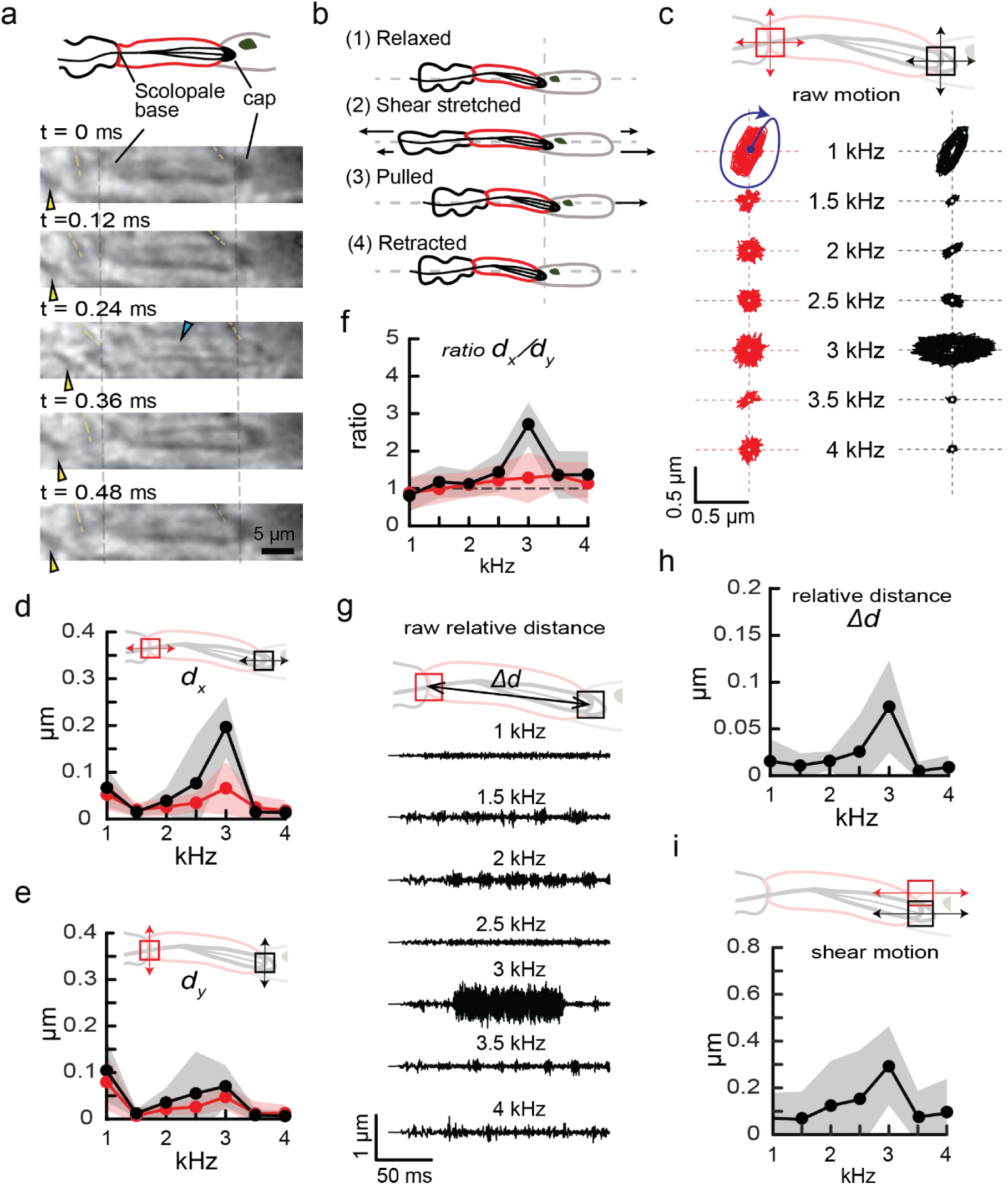
Sound-evoked motion of the scolopidium. **(a)** Schematic of a scolopidium showing the scolopale base and the cap that are clearly visible in light microscopy. (top). Montage over a period of one cycle of sound-evoked motion of a Group-III scolopidium stimulated with 3 kHz pure tone at 90 dB SPL (bottom). At t = 0.24 ms, the cyan arrowhead indicates the cilium when it is brought into focus. The yellow lines indicate the shear near the cap and the scolopale base. The yellow arrowheads follow the region with clear deformation near the scolopale base. **(b)** Summary of the sound-evoked motion of the scolopidium. **(c)** Motion of the scolopale base (red) and cap (black) regions along (dx) and orthogonal (dy) to the ciliary axis. **(d)** Root-mean-square (RMS) amplitude of the scolopale base (red, N = 6, n = 22) and the cap (black, N = 6, n = 22) regions along the ciliary axis (dx). **(e)** RMS amplitude of the scolopale base (red, N = 6, n = 22) and the cap (black, N = 6, n = 22) regions along the orthogonal axis (dy). **(f)** Ratio of the axial-orthogonal motion (dx/dy) of the scolopale base (red, N = 6, n = 22) and the cap (black, N = 6, n = 22). **(g)** Example of distance (Δ𝑑) between the scolopale base and the cap region as a function of time over each stimulated frequency. **(h)** Relative distance between the scolopale base and the cap (N = 6, n = 24). **(i)** Shear motion along the ciliary axis at the cap region (N = 6, n = 22. For additional data, see Extended Data Fig. 3 and Source Data Fig. 4.

First, we analysed the raw amplitude of the scolopale base and the cap. Upon sound stimulation, the entire scolopidium underwent elliptical motion (Fig. 4c). Notably, the motion of the scolopale base was generally more circular, while that of the cap was more elliptical, particularly between 1.5-3 kHz. At 1 kHz; however, the scolopidium showed a strong elliptical motion oblique to the ciliary axis. When stimulated at 3 kHz, the elliptical motion of the cap aligned with the ciliary axis and was most pronounced.

To investigate the principal axis of the sound-evoked motion, we decomposed the motion of both the cap and the scolopale base into components along the ciliary axis, 𝑑_𝑥_, (Fig. 4d) and orthogonal to it, 𝑑_𝑦_, (Fig. 4e). For both landmarks, motion along the ciliary axis was strongest, with a prominent peak at 3 kHz, base 𝑑_𝑥_: 66 ± 55 nm (N = 6, n = 24) and cap 𝑑_𝑦_: 197 ± 64 nm (N = 6, n = 24), (Fig. 4d, Extended Data Fig. 3a, c). In contrast, orthogonal motion was minimal, with only a shallow peak at the same frequency, base 𝑑_𝑥_: 47 ± 26 nm (N = 6, n = 24) and cap 𝑑_𝑦_: 70 ± 44 nm (N = 6, n = 24), (Fig. 4e, Extended Data Fig. 3b, d). Consequently, the ratio of axial-to-orthogonal motion, 𝑑_𝑥_/𝑑_𝑦_, is consistent with elliptical motion shown in Fig. 4c, where cap 𝑑_𝑥_/𝑑_𝑦_ peaked sharply at 3 kHz, base 𝑑_𝑥_/𝑑_𝑦_: 1.29 ± 0.66 (N = 6, n = 24) and cap 𝑑_𝑥_/𝑑_𝑦_: 2.72 ± 1.00 (N = 6, n = 24), confirming that axial motion is the dominant component of the frequency response (Fig. 4f).

Since the adequate stimulus for MET is expected to arise from relative motion within the scolopidium, we next computed (i) the relative distance between the cap and the scolopale base, Δ𝑑, and (ii) shear motion at the cap region quantified by tracking the upper and lower borders of the attachment cell upon sound stimulation. Indeed, the relative motion was also strongest and sharply peaked at 3 kHz, Δ𝑑: 74 ± 48 nm (N = 6, n = 24) (Fig. 4g-h). The shear motion of the cap region; however, produced a distinct but broad peak around 3 kHz, 215 ± 200 nm (N = 6, n = 22) (Fig. 4i).

### 2.5. Direct pulling of the scolopidium cap evokes maximal transduction current

High-speed light microscopy revealed that scolopidial motion during sound stimulation occurred primarily along the ciliary axis, with additional components orthogonal to it (Fig. 4). To identify the principal axis through which the scolopidium is stimulated, we directly mechanically stimulated the scolopidium cap with a stimulating pipette while measuring transduction currents via whole-cell patch-clamp recordings of the corresponding Group-III auditory neuron somata (Fig. 5a; Supplementary Video 3). We delivered a series of step displacements in two directions: a ’pull-push’ (horizontal) motion along the ciliary axis (Fig. 5b) and an ’up-down’ (vertical) motion orthogonal to it (Fig. 5c). Here, we showed that stimuli in all four directions evoked sharp, transient inward currents at the onset of the step (Fig. 5d-e). The most sensitive responses were to pull and push stimulations at 1 nm (Fig. 5d), whereas upward and downward stimulations produced clearly noticeable transduction currents from 5 nm and above (Fig. 5e). The signal-to-noise ratio in single trials was low due to spontaneous discrete depolarisations found in chordotonal organs^7,9,33^. Upon averaging the responses over the total number of locusts, pull-push: N = 18, n = 18 and up-down: N = 11, n = 11, the ensemble average transduction current revealed a graded, non-linear relationship between stimulus and current (Fig. 5f-g).

**FIGURE 5.**
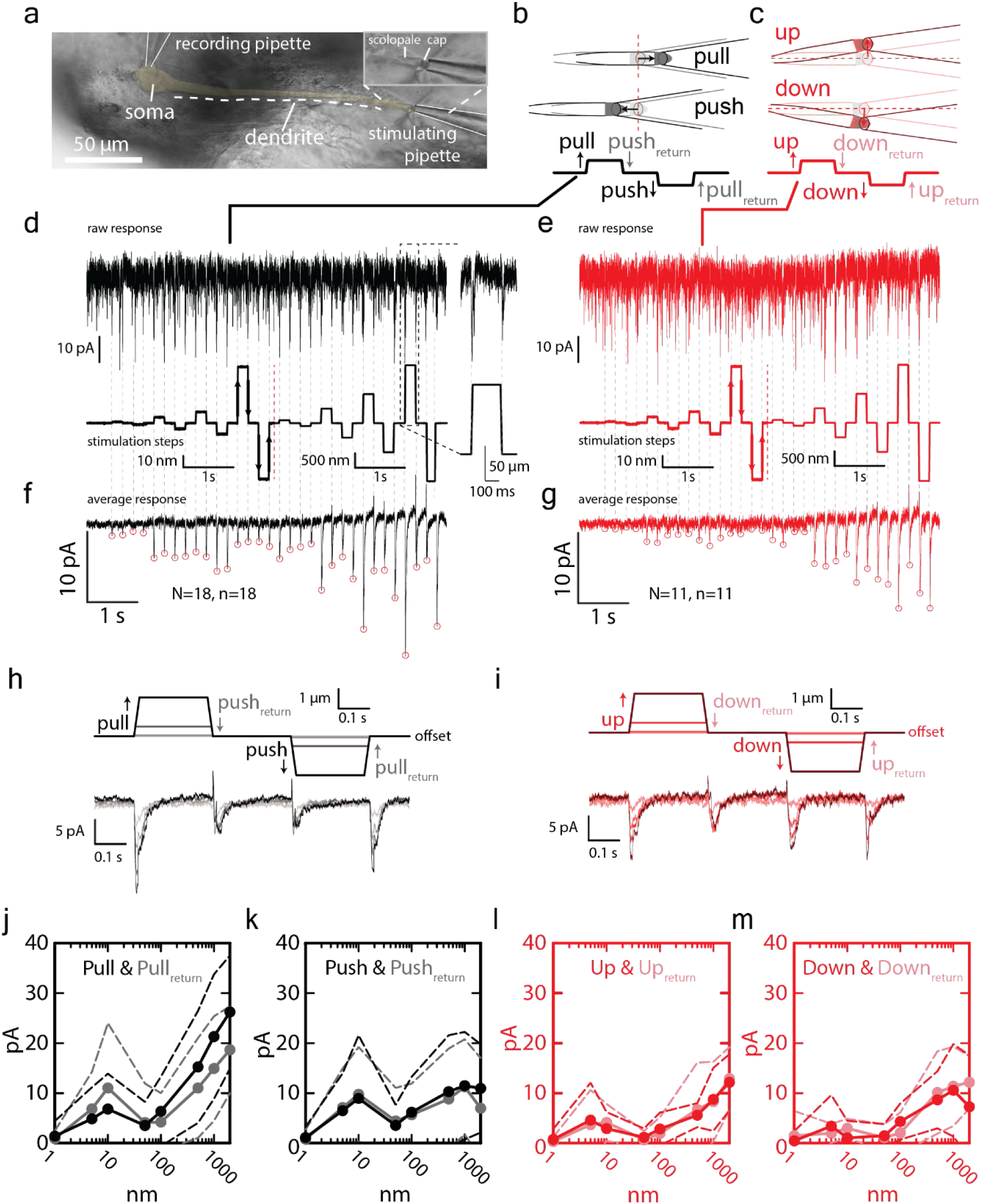
Transduction current upon step displacements of a scolopidium cap. **(a)** Focal-stacked light microscopy image from ∼10 µm apart showing the auditory neuron (yellow), a patch recording pipette, and a stimulating pipette. **(b-c)** Schematics of step-displacement protocols for (b) pull-push displacements along the ciliary axis (N = 18, n = 18) and (c) up-down displacement orthogonal to the ciliary axis (N = 11, n = 11). **(d-e)** Transduction currents (top trace) in response to (d) pull-push and (e) up-down step-displacements with increasing amplitudes (bottom trace). Step-displacement traces below and over 100 nm in (d-e) are shown in different scales. Arrows indicate the orientation of stimulation. **(f-g)** Ensemble average transduction currents in response to (f) pull-push and (g) up-down step displacements. The stimulation traces are shown in (d-e). **(h-i)** Examples of the transduction currents in response to (h) pull-push and (i) up-down step displacement of magnitudes 2, 50, 500, and 2000 nm. **(j-m)** Ensemble average transduction current plotted as a logarithmic function of the step-displacement, I-log(X) curve, for (j) pull, (k) push, (l), and (m) downward displacements with their return to the offset counterparts. For additional data, see Source Data Fig. 3.

Next, we quantified transduction currents in response to pull-push step stimuli averaged across all recordings. There was a general increase in the magnitude of the transduction current with increased step displacement (Fig. 5h, i). Discrete depolarisations contributed to a large standard deviation of step-evoked transduction currents (Fig. 5d, e). There were distinct bi-phasic and logarithmic phases in the current-displacement relationship, I-log(X) curve, from 1-10 nm and 50-2000 nm in both pull-push (Fig. 5j, k) and up-down (Fig. 5l, m) displacements. The logarithmic relationship was determined by fitting linear regression models to the I-log(X) curves (see Source Data Fig. 5). The largest transduction currents were produced by pull along the ciliary axis. We quantified the difference at 2,000 nm displacement from the resting position between pull, 26.22 ± 10.99 pA (N = 18, n = 18), and push, 10.96 ± 8.59 pA (N = 18, n = 18) (Tukey’s procedure, *p* = 3.1e-05, t = -9.58). Next, we found that the transduction current was larger for pull_return,_ 18.68 ± 8.43 pA (N = 18, n = 18) compared to push_return_ 6.99 ± 9.75 pA (N = 18, n = 18) (Tukey’s procedure, p = 0.003, t = -4.72). Additionally, the transduction current evoked by pull from the resting position was larger than displacement upward, 12.17 ± 5.37 pA (N = 11, n = 11) (Tukey’s procedure, *p* = 1.75e-03, t = -3.83) or downward 7.28 ± 9.76 pA (N = 11, n = 11) (Tukey’s procedure, *p* = 5.10e-06, t = -4.53). There was no difference between pull_return_ and up_return_ (Tukey’s procedure, *p* = 0.69, t = -1.91), and between pull_return_ and down_return_ (Tukey’s procedure, *p* = 0.55, t = -2.24).

## 3. Discussion

The scolopidium is the basic sensory unit of mechano-electrical transduction (MET) in chordotonal organ of insects^3,10,12,13,17,27^. Addressing the fundamental question of how the scolopidium operates, we demonstrate here that axial stretch along the cilium is the adequate stimulus or the key mechanical input that activates mechano-electrical transduction. This conclusion is based on two key experiments. First, high-speed imaging revealed that elliptical sound-evoked scolopidium motion is dominated by stretch along the ciliary axis (Fig. 4). Second, pull of the scolopidium along the ciliary axis evoked the maximal MET current (Fig. 5).

Axial stretch is assumed to be the adequate stimulus in other chordotonal organs, based on either ultrastructural analysis or electrophysiological recordings, but this has never been experimentally confirmed with direct observation of the underlying mechanics^10,12,34^. These early studies, performed before the molecular identity and precise location of MET channel candidates like NompC and Nan-Iav were known, led to at least ten proposed mechanisms^5,35^. For example, deformation of the ciliary dilation^10,12,34^, bending of the cilium^12,34^ or scolopale sheath^25,36^, and contraction of the ciliary rootlet ^36^. Yet another hypothesis proposed that the cap that encloses the cilium could narrow during axial stretch and exert a compressive force to the ciliary membrane^13^. Our finding provides the first direct evidence that axial stretch is the adequate stimulus of this highly conserved mechanoreceptor found in all insects and some crustaceans^37^.

How does stretch of the scolopidium lead to conformational change of MET channels within the cilium? Here, we propose four competing hypotheses (Fig. 6): (i) In the *‘direct-stretch model’*, axial tension is transmitted along the ciliary membrane, stretches the lipid bilayer and opens MET channels. (ii) In the *‘stretch-compression model’*, axial stretch narrows the central passage of the cap thus compressing the ciliary tip i.e. when a compressive material with a hollow tube is deformed along its axis, the tube narrows (like a ‘Chinese finger trap’). (iii) In the *‘stretch-deformation model’*, the larger ciliary dilation is pulled against and into the narrow central passage of the cap (Fig. 2e, f). (iv) In the *‘stretch-tilt model’,* the cap pivots due to the shear component of the scolopidium stretching, this pivoting leads to either asymmetrical axial tension along the ciliary membrane or asymmetrical compression of the narrow central passage against the ciliary tip. In short, the proposed models differ in the primary force that opens the MET channels. The MET channels would be opened either through membrane tension in the direct-stretch model, or by compressive forces of the cap wall directly onto the membrane in the stretch-compression and stretch-deformation models. However, in the stretch-tilt model the MET channels could be activated through either membrane tension as the cilium bends, lateral compression as it presses against the cap wall, or a combination of both.

**FIGURE 6.**
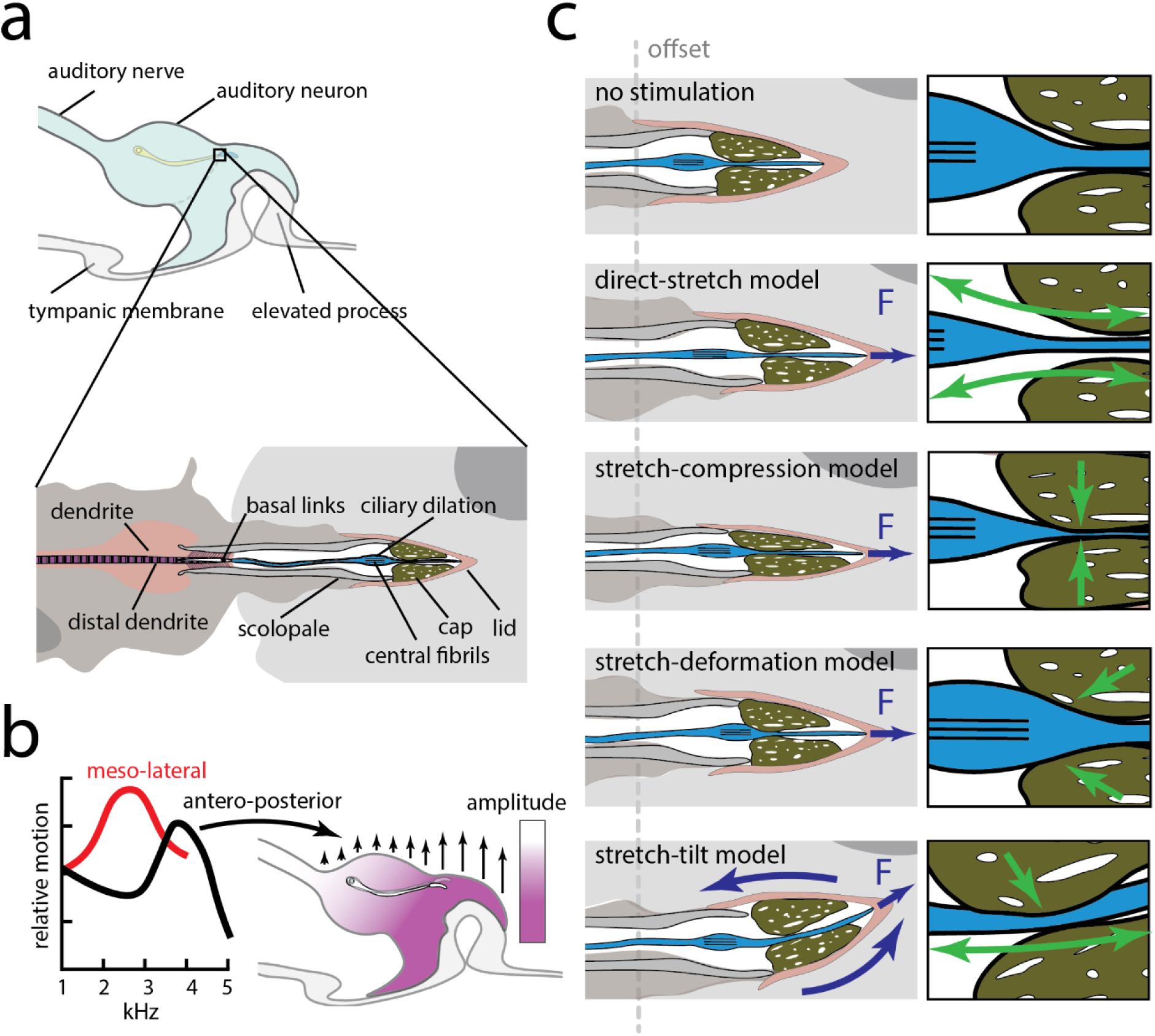
Possible gating mechanisms of MET channels in insect chordotonal neurons. **(a)** Key anatomical features of Müller’s organ and the scolopidium. **(b)** Sound-evoked motion of Müller’s organ generates axial stretch at the level of the scolopidium. **(c)** Possible gating mechanisms of the mechano-electrical transduction channels in insect chordotonal neurons (left): (i) direct-stretch model, (ii) stretch-compression model, (iii) stretch-deformation model, and (iv) stretch-tilt model. Purple arrows indicate forces exerting on the scolopidium. The right panel shows magnified views of the entry of the cap’s central passage, showing forces (green arrows) exerting on the ciliary membrane.

While a complete gating model for insect MET remains elusive^38^, key molecular insights have been provided by the cryo-EM structure of the putative channel NompC^39^, as well as by Molecular Dynamics (MD) simulations^40,41^, and electrophysiological recordings^40,42,43^. Specific structures within NompC, such as its Ankyrin Repeat (AR)^43,44^ and Linker Helix (LH)^42^ domains, have been proposed to form a “gating spring”—a compliant element that transmits mechanical force into the channel’s conformational opening^42,45^.

Currently, there is no direct evidence to support the direct-stretch model. However, there are lines of compelling evidence that support the stretch-compression, stretch-deformation, and stretch-tilt models. In particular, NompC expressed in *Drosophila* S2 transfected cells are more sensitive to membrane compression than suction^40^. The other MET channel candidate Nan-Iav is not mechanosensitive when expressed in HEK293T cells^8^. The properties of both of these Transient Receptor Potential family ion channels may operate differently in the cilium compared to heterologous cell lines. Nanchung-Inactive is expressed below the ciliary dilation in the proximal ciliary segment. Therefore, if Nanchung-Inactive ion channels were mechanically sensitive *in vivo* they could only be opened by stretch as the stretch-compression, stretch-deformation and stretch-tilt models act only on the ciliary tip. Our OCT-based vibrometry reveals that axial stretch of the scolopidium arises from complex 3D motion of Müller’s organ^21^, a feature that maybe shared by other chordotonal organs^14,46,47^. We demonstrate an abrupt transition in both phase and amplitude localised at the site of Group-III scolopidia, which would generate relative motion across the ciliary axis (Fig. 3 g, h). While motion along the antero-posterior axis is largely dominated by the tympanum’s movement at 3.5-4 kHz^29^, strong vibrations along the meso-lateral axis at lower frequencies, 2.25-3 kHz (Fig. 3c, e), may contribute to broad frequency sensitivity of the sound-evoked transduction current^1^ . In this experiment, we measured vibration along the two axes independently. To fully capture complex 3D motion of Müller’s organ, simultaneous measurement of vibration in both axes warrants further investigation, using either a two-OCT recording beam setup or a mirror to reflect the OCT beam onto an additional axis^48^.

Finally, our high-resolution 3D ultrastructure provides the anatomical framework for how the scolopidium operates (Fig. 2). The presence of the scolopale ’lid’ and the full extent of the cilium to its insertion at the scolopale cartridge tip indicate a firm, direct coupling between the attachment cell and the cilium, essential for a strong anchor at the cilium’s distal end. Furthermore, the distinct narrowing at the cilium’s entrance to the central passage may serve to compress or deform the cilium during axial stretch (Fig. 2e, f). The numerous oblique basal links may not be directly involved in transduction but likely provide mechanical stability to the distal dendrite and ciliary rootlet (Fig. 2h, i). The nature of the central fibrils within the ciliary dilation (Fig. 2j, k) remains unknown but are also in bush cricket and mosquito auditory neuron cilia^14,49^. Our OCT and high-speed camera measurements are from *ex vivo* preparations immersed in saline solution and our transduction current recordings are from enzymatically dissociated scolopidia. Despite these manipulations, the electrophysiological function of the auditory neurons appears physiologically normal when compared against preparations that more closely mimic *in vivo* conditions^4,9,14,33,50^. In our previous work, we showed that, Group-III neurons retain their frequency tuning even when Müller’s organ and the internal surface of the tympanum is bathed in saline^1^. Thus, we conclude that the electrophysiology of the auditory neurons and the mechanics of Müller’s organ and the scolopidia reported here mimic *in vivo* conditions.

In conclusion, our study provides a comprehensive, multi-scale analysis of MET in an insect auditory chordotonal neuron, using excised preparations of locust ears. We have shown that complex 3D motion of Müller’s organ produces localised relative motion precisely where the auditory scolopidia are situated. This relative motion within the organ then drives an axial stretch of the scolopidium, which in turn evokes the mechano-electrical transduction current. As scolopidial structure is ubiquitous across insect classes, we find it probable that one of the four mechanisms of MET channel opening that we proposed here is largely conserved across insects.

## Acknowledgments

We thank Daisy Ogle and Alix Blockley for providing z-stack images of Dylight-488 conjugated streptavidin neurobiotin labelled auditory neurons, which allowed us to map the positions of Group-III scolopidia. We are grateful to Celia Hansen and Ramesh Patel for maintaining the locust facility. Deep appreciation to Anna Vavakou who assisted us in the initial Optical Coherence Tomography (OCT) experiments. We thank the following scientists who helped us with sample preparation and data acquisition. Focused Ion Beam Scanning Electron Microscopy (FIB-SEM) was performed by Claire Boulogne at the Electron Microscopy facility of Imagerie-Gif, Institute for Integrative Biology of the Cell (I2BC). Sample preparation for electron microscopy was assisted by Natalie Allcock and Josh Whittingham. An X-ray micro-CT scan image was performed by Gareth Douglas. We also thank Kanpong Boonthaworn, Mei-ling Joiner, and Daniel Eberl for their valuable feedback on the manuscript. This project was funded by a Long-term Human Frontier Science Program (HFSP) Fellowship (LT0002/2023-L) and a France Bioimaging Access Grant (Access-4942) awarded to A.C., as well as a University Research Fellowship from the Royal Society awarded to B.W. (URF\R1\180022 & URF\R\231029).

## 4. Author contributions

A.C. and B.W. designed and planned the experiments. A.C. performed sample preparation, acquisition, annotation, and three-dimensional reconstruction for Focus-Ion Beam Scanning Electron Microscopy (FIB-SEM). M.V.H. and M.N. performed the initial Optical Coherence Tomography (OCT) experiments, A.C. and B.W. finalised the data collection and analysis. B.W. performed whole-cell patch clamp recordings. A.C. and B.W. analysed the experimental data and prepared the initial version of the manuscript. M.N. and M.V.H. commented on it, and together with A.C. and B.W., finalised the submitted version of the manuscript.

## EXTENDED DATA FIGURES

**EXTENDED DATA FIGURE 1.**
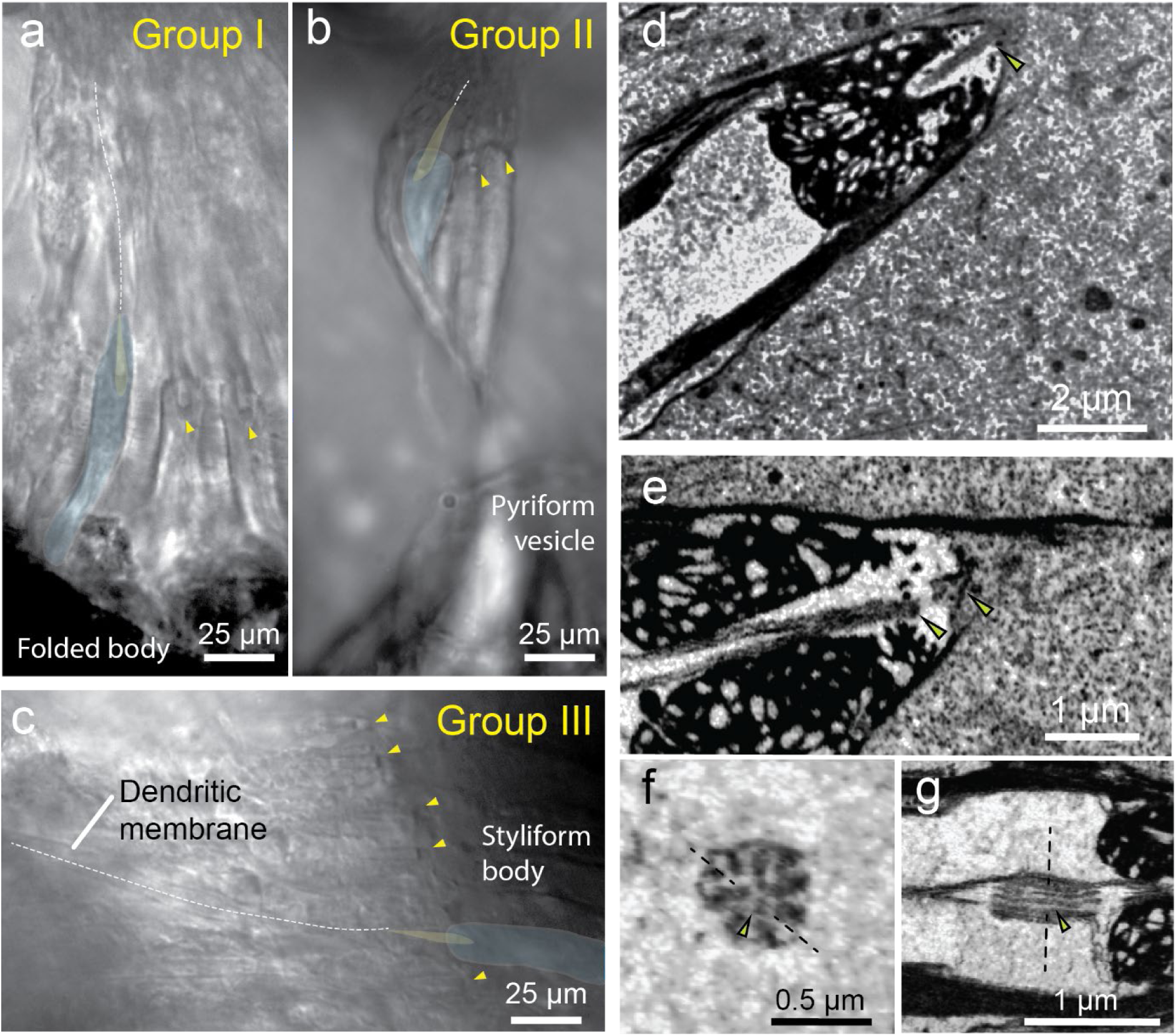
Anatomy of the scolopidia. **(a-c)** Locations of the three groups of scolopidia in Müller’s organ: Group-I scolopidia, located near the folded body; Group-II, located within the fusiform body, forming a ’bridge’ to the pyriform vesicle at the surface of the tympanic membrane; and Group-III, located on the styliform body. The overlay indicates examples of scolopale sheaths (opaque yellow) and the corresponding attachment cells (blue). Dashed lines outline the dendrites along their lengths, and yellow arrowheads mark the other visible scolopidia. **(d)** Oblique section of the distal region of a scolopidium reveals the full extent of the cilium. **(e)** Example of a potential sample preparation artifact showing the apparent detachment of a cilium from its terminal region. **(f)** Longitudinal section through the ciliary dilation, revealing three central fibrils (arrowhead). **(g)** Longitudinal section along the dashed line in (f), reveals two of the central fibrils (arrowhead) extending along the entire length of the dilation.

**EXTENDED DATA FIGURE 2.**
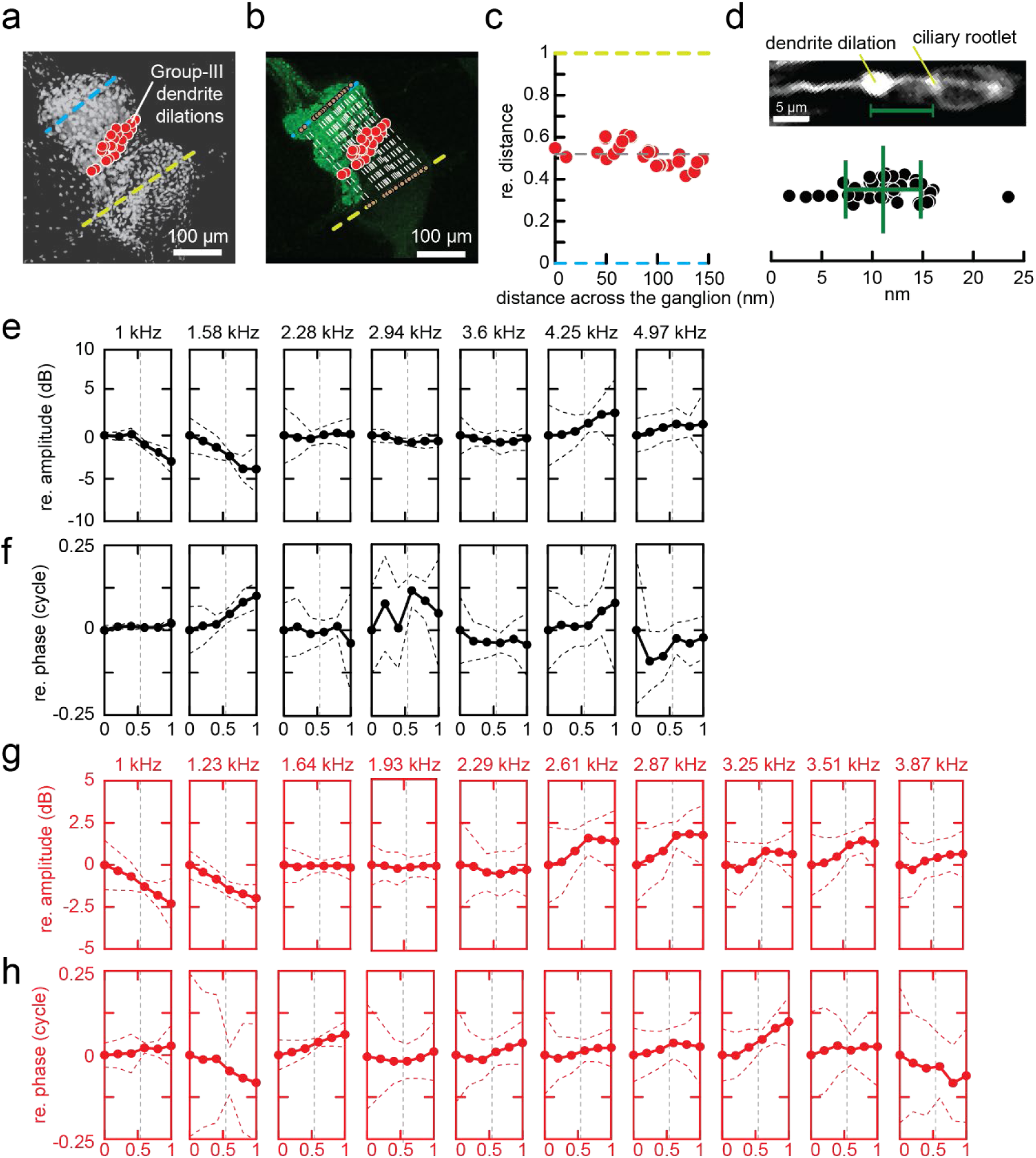
Locations of Group-III scolopidia and frequency response of Müller’s organ using OCT. **(a)** Maximal intensity projection of a Müller’s organ labelled with DAPI superimposed with manual registration of dendrite dilations (red circles). Anatomical landmarks are indicated at the maximal width of the ganglion (cyan dashed line) and the styliform body (yellow dashed line). **(b)** Maximal intensity projection of auditory neurons labelled with DyLight™ 488-conjugated streptavidin neurobiotin superimposed with manual measurement of dendrite dilations’ positions (red circles) and schematic illustrating the computation of the relative distance of each dendrite dilation between the ganglion (cyan dashed line) and the styliform body (yellow dashed line). **(c)** Normalised positions of dendrite dilation from an example of one Müller’s organ. The normalised positions of the dendrite dilations are plotted as a function of distance across the ganglion’s width shown as cyan dashed line in (b). A black dashed line indicates the normalised position at 52% between the ganglion and the styliform body. **(d)** Distances between dendrite dilations and the ciliary rootlet of the corresponding scolopidium (N = 19, n = 46). **(e, f)** Amplitude (e) and phase (f) response of Müller’s organ along the antero-posterior axis (N = 4, E = 4) measured using OCT, plotted as a function of relative distance between the ganglion and styliform body. **(g, h)** Amplitude (g) and phase (h) response of Müller’s organ along the meso-lateral axis (N = 7, E = 12) measured using OCT, plotted as a function of relative distance between the ganglion and styliform body. The vertical dashed lines in (e-h) indicate the normalized position of the Group-III scolopidia. All vibration amplitudes are shown in dB relative to a 1-nm reference vibration. For additional data, see Source Data Extended Fig. 2.

**EXTENDED DATA FIGURE 3.**
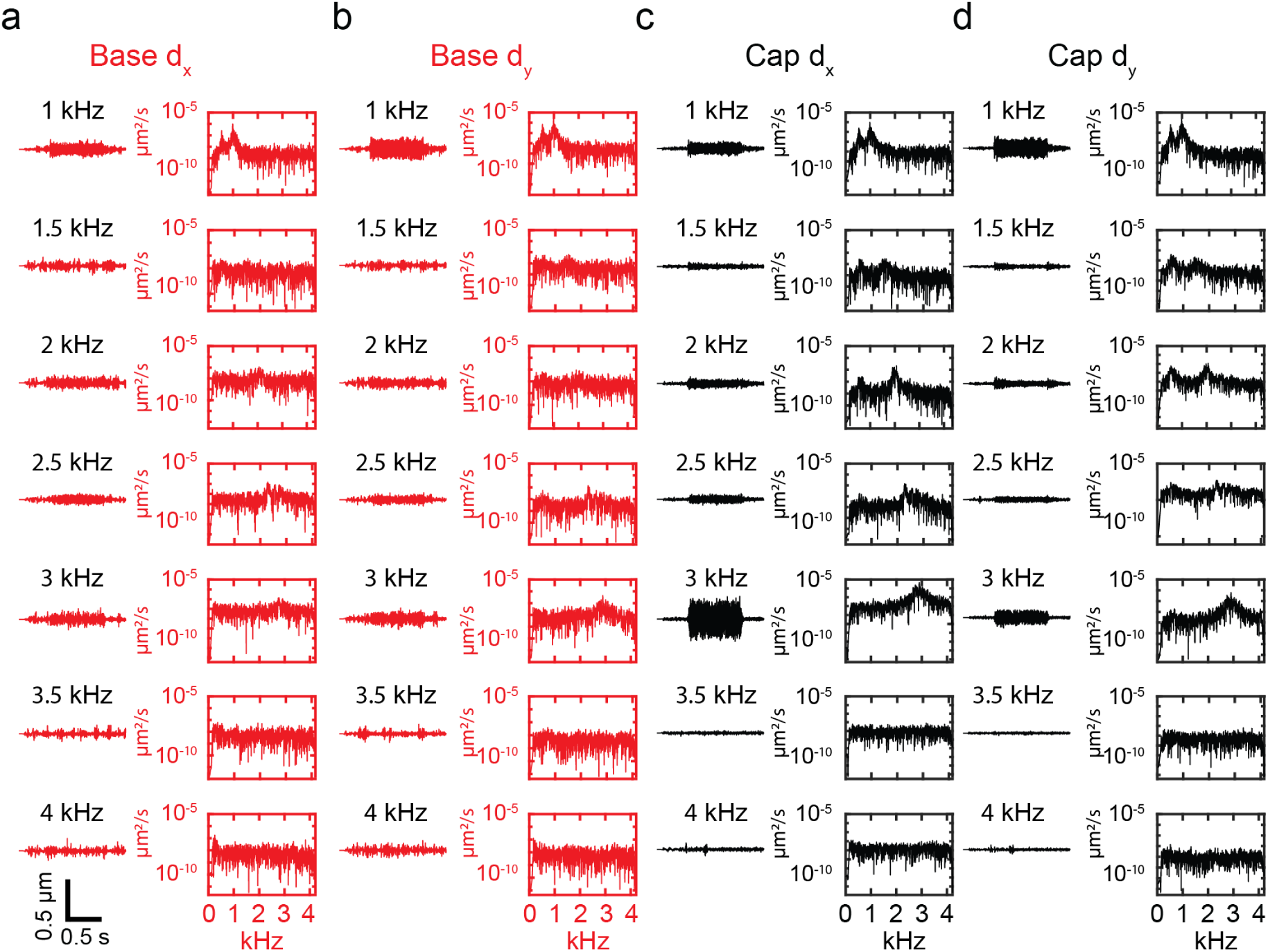
Example recordings of sound-evoked motion of a scolopidium. Raw time series amplitude traces (left traces) and their corresponding Fourier transforms (right traces) for the scolopale base and cap along (dx) and orthogonal (dy) to the ciliary axis at each stimulated frequency from 1 to 4 kHz with an increment of 0.5 kHz. **(a, b)** Motion of the scolopale base (a) along and (b) orthogonal to the ciliary axis. **(c, d)** Motion of the cap (c) along and (d) orthogonal to the ciliary axis. Pure-tone sound stimulation at 90 dB SPL.

## METHODS

### Animals and sample preparation

Desert locusts, *Schistocerca gregaria*, were reared under crowded conditions with a 12:12-hour light/dark cycle and the ambient temperature at 36 °C (light) and 25 °C (dark). They were fed fresh wheat and bran. All experiments used adult male and female locusts 10-20 days after their final moult.

We used an excised locust ear preparation for all experiments, following a previously described dissection protocol^9^. In short, the ear was surgically removed and affixed to a custom-made 3D-printed plastic dish, positioning the tympanum over a 3 mm hole. For all measurements, the external side of the tympanum faced the sound source, while the internal side, including Müller’s organ, was submerged in locust extracellular saline. The extracellular saline contained 185 mM NaCl, 10 mM KCl, 2 mM MgCl_2_, 2 mM CaCl_2_, 10 mM HEPES, 10 mM trehalose, and 10 mM glucose. The pH was adjusted to 7.2 using NaOH.

### Sample preparation for FIB-SEM and 3D reconstruction of a scolopidium

Excised locust ears were fixed in a solution of 2.5% glutaraldehyde and 2% paraformaldehyde in 0.1 M sodium cacodylate (pH 7.2) with 4.1% sucrose. After washing, the samples were incubated in 1% osmium tetroxide with 1.5% potassium ferricyanide, followed by another wash and a 20-minute incubation in thiocarbohydrazide. Samples were then washed and sequentially incubated in 2% aqueous osmium tetroxide and 2% uranyl acetate, with washing steps in between. Following fixation and staining, the samples underwent serial dehydration in ethanol (30%, 50%, 70%, 90%, and 100%). The samples were then washed in propylene oxide before being embedded in a mixture of hard-formula TAAB 812 resin and propylene oxide. The resulting resin blocks were trimmed to the region of interest on an ultramicrotome (Leica UC7), which was verified by staining 100 nm-thick sections with methylene blue.

Serial acquisition was performed on a dual-beam FIB-SEM (Zeiss Crossbeam 500) equipped with a Gemini2 SEM column and a Ga^+^ source. Imaging parameters were set as follows: SEM extra high tension (EHT) at 1.6 kV with a 600 pA beam current, and FIB current at 700 pA at 30 kV. The voxel sizes of the acquired FIB-SEM stacks ranged from 9 to 15 nm. The 3D ultrastructure of the scolopidia were reconstructed from the FIB-SEM stacks using image processing software Dragonfly 3D.

### Optical Coherent Tomography Vibrometry

We measured sound-evoked vibrations using a spectral domain Optical Coherence Tomography (OCT) system (Ganymede, Thorlabs). The field of view was 10 × 10 mm with a depth range of 3.5 mm. The OCT beam was directed from above, while the preparation was mounted either horizontally (with the sound source below, Fig. 3a) or vertically (with the sound source to the side, Fig. 3b). The excised ears were submerged in large amount of saline solution (∼10 mm liquid high above the surface of the tympanic membrane) to minimise drifting over the time course of acquisition due to evaporation. The beam was manually aligned for each measurement axis to maximize the signal-to-noise ratio for that orientation. Sound stimuli, consisting of broadband multitone ‘zwuis’ complexes^51^, were generated using a Tucker-Davis Technologies System III and delivered via a ScanSpeaker R2904/700 005 loudspeaker, placed at 32 cm from the preparation. The multitone ‘zwuis’ method allows for measurement of vibration amplitudes and phases from several frequencies simultaneously. The stimulus level used was 90 dB SPL; based on previous work, this level in a liquid-backed preparation corresponds to approximately 70 dB SPL in an air-backed tympanum^52^.

OCT acquisitions were externally triggered and phase-locked to the acoustic stimulation at a sampling rate of 90 kHz. To characterize the organ’s complex 3D motion, vibrations were measured separately along two primary axes: antero-posterior (motion normal to the tympanic surface) and meso-lateral (tangential, side-to-side motion). The vibration magnitude is heavily damped but comparable to those submerged in a similar level of saline solution and measured using Laser Doppler Vibrometer (LDV)^52^. Vibration waveforms were derived from contiguous groups of three pixels per scan. All organ vibrations were measured relative 1 nm. The significance of phase-locking for each frequency component was assessed using the Rayleigh test (p = 0.001). All data acquisition and analysis were performed using a custom MATLAB toolbox (MathWorks, 2007) developed by Marcel van der Heijden.

The vector differences (Fig. 3e) were computed from the complex amplitudes of the two vibrations. Each vibration was represented as a complex number, 𝐴𝐴 · 𝑒𝑒^𝑖𝑖𝑖𝑖^, where 𝐴𝐴 is the vibration amplitude and 𝜑𝜑 is its phase relative to the stimulus. The vector difference represents the magnitude of the resulting complex number after subtracting the complex amplitude at the styliform body from that at the ganglion.

### Localisation of Group-III scolopidia

To prepare the locusts for neural tracing, each insect was secured on its back in plasticine. A small section of the cuticle on the second and third thoracic segments was carefully cut and temporarily removed. After clearing away the tracheal air sacs to expose the metathoracic ganglia and the auditory nerve (nerve VI), the thoracic cavity was filled with a saline solution. The auditory nerve was then severed near the ganglion, and the cut end was positioned on the outer cuticle at the rear of the thorax. A small well was made around the exposed nerve ending using petroleum jelly and filled with a 5% Neurobiotin solution to allow the tracer to diffuse along the nerve. The well was sealed with more petroleum jelly, and the previously removed cuticle was placed back onto the original area and also sealed to prevent the area from drying out. The locusts were then incubated overnight at 4°C to facilitate the diffusion of Neurobiotin along the auditory neurons. After the incubation, the locust ears were dissected from the first abdominal segment. These ears were fixed in a 4% paraformaldehyde solution for 24 hours at 4°C. Subsequently, the fixed ears were washed three times in Phosphate Buffer Saline (PBS). To prepare for imaging, the ears were treated to make the cells permeable and to block non-specific binding. They were then incubated overnight with Dylight 488 streptavidin, which binds to the Neurobiotin to stain the auditory neurons, and DAPI, which stains the cell nuclei. Finally, after another series of washes in PBS, the ears were dehydrated using an ethanol series and cleared with Methyl salicylate. The stained neurons were then imaged using a confocal microscope.

The positions of dendrite dilations were then manually measured in Fiji^53^. The positions were recorded and saved as the region of interest (ROI) files and further processed in MATLAB R2004a (Mathworks). The position of each dendrite dilation was computed as a relative distance between the ganglion and the styliform body (Extended Data Fig. 2).

### Tracking sound-evoked scolopidium motion

We recorded the sound-evoked motion of the scolopidium using a high-speed camera (KINETIK, Teledyne) mounted on an FN1 Nikon upright microscope using a 40X/0.8W WD = 3.5 mm (NIR Apo DIC N2) objective. Using an objective with a small working distance enabled us to minimise the saline volume, which reduced damping and maximised the amplitudes of scolopidial motion within the detection range of our imaging system. We achieved an average acquisition rate of 8,602 ± 289 frame per second (24 movies). Sampling with an acquisition rate that did not achieve Nyquist sampling of its stimulated frequency were excluded from analysis. Sound stimulation was synchronised with imaging and delivered through a speaker (RS PRO 50 mm, RS PRO) via an analogue-to-digital converter (National Instruments PCIe 6323). The sound pressure level was calibrated by measuring the voltage output of an amplified microphone (Clio PRE-01 and Clio MIC-03, Audiomatica) in response to a 94 dB SPL 1 kHz tone from a sound level calibrator (Cal73, Brüel & Kjær). This calibration allowed us to determine the sound pressure level from the microphone’s voltage output. The microphone was then positioned where the excised locust ear would be during experiments, and speaker voltages were adjusted to produce a flat response across the stimulated frequency range (1–5 kHz). An in-house MATLAB graphical user interface controlled the setup. In these experiments, we stained the excised locust ear preparation with 0.1% Janus Green B (201677-5G, Sigma Aldrich) to enhance the contrast of the scolopidium with its surrounding tissue.

The sub-pixel resolution motion of the scolopidium was tracked using Fiji/ImageJ Template Matching plugin (https://sites.google.com/site/qingzongtseng/template-matching-ij-plugin)^54^. The tracking method is based on Normalised Correlation Coefficient and Gaussian peak fitting to determine the location of the correlation peak. The tracking achieved sub-pixel resolution of tens of nanometres. The coordinates output from the plugin were further analysed in MATLAB R2004a (Mathworks).

### Whole-cell patch clamp and mechanical stimulation of a scolopidium

To expose Group-III auditory neurons for patch-clamp recordings, the fibrous tissue enclosing the medial–dorsal border of Müller’s organ was digested by applying a solution of collagenase (0.5 mg/ml) and hyaluronidase (0.5 mg/ml; C5138 & H2126, Sigma-Aldrich) in extracellular saline. The enzymes were applied locally using a wide-bore (12 µm) patch pipette, which was also used to gently remove the softened tissue via suction, exposing the somatic membrane of the neurons.

To prepare for mechanical stimulation, the cap of a targeted scolopidium was released from its attachment cell by puffing the same enzyme solution from the same wide-bore pipette. A new pipette filled with extracellular saline was then used to hold the cap via gentle, continuous mouth suction, which was renewed approximately every minute, and before each recording series, to ensure a secure connection. A series of push-pull (axial) and up-down (normal) step displacements (1, 5, 10, 50, 100, 500, 1000, and 2000 nm) were delivered to the stimulating pipette using a PatchStar micromanipulator (Scientifica).

Scolopidia were visualized with a custom upright microscope (Cerna® Mini, Thor Labs) equipped with a 40× water-immersion objective (0.8 NA, 2 mm working distance; CFI APO NIR, Nikon). Patch-clamp electrodes (3-4 MΩ) were fabricated from borosilicate glass (GB150–8P, Science Products) using a vertical pipette puller (PC-100, Narishige). The intracellular (pipette) solution contained (in mM): 170 K-aspartate, 4 NaCl, 2 MgCl_2_, 1 CaCl_2_, 10 HEPES, 10 EGTA, and 20 tetraethylammonium (TEA). The extracellular saline contained (in mM): 185 NaCl, 10 KCl, 2 MgCl_2_, 2 CaCl_2_, 10 HEPES, 10 trehalose, 10 glucose, and 0.09 tetrodotoxin (TTX) citrate. To isolate and increase the magnitude of the transduction current, spikes were blocked with extracellular TTX and intracellular TEA. Electrophysiological signals were amplified, digitized, and recorded using an EPC 10 USB patch-clamp amplifier (HEKA Elektronik).

The data analysis was done in MATLAB R2004a (Mathworks). Peak detections of the ensemble average transduction current (Fig. 5 f, g) were done by detecting peak within a 30-ms windows at the onset of step-stimulation using Matlab built-in peak detection package ‘findpeaks’.

### Statistical analysis

Descriptive data are presented as mean ± standard deviation. Biological replicates are denoted by N (number of locusts), E (number of ears), and n (number of cells). For the statistical analysis in Fig. 5j-m, we fitted linear regression models to assess the correlation between the transduction current and the logarithmic step displacement from 50-2000 nm. To test for statistical differences, we performed a one-way ANOVA followed by Tukey’s Honestly Significant Difference (HSD) post-hoc test for multiple comparisons. The reported p-values were obtained from the Tukey HSD procedure, while the t-statistics are from a Student’s t-test. Further details on the statistical analysis are available in the source data.

### Data availability

Raw data and MATLAB codes supporting plots in this manuscript can be found in the online repository Zenodo^55^. Source data for each figure are provided with this paper as stated in the figure legends. Additional data can be requested from the corresponding author upon reasonable request.

## SUPPLEMENTARY INFORMATION

**SUPPLEMENTARY MOVIE 1.**
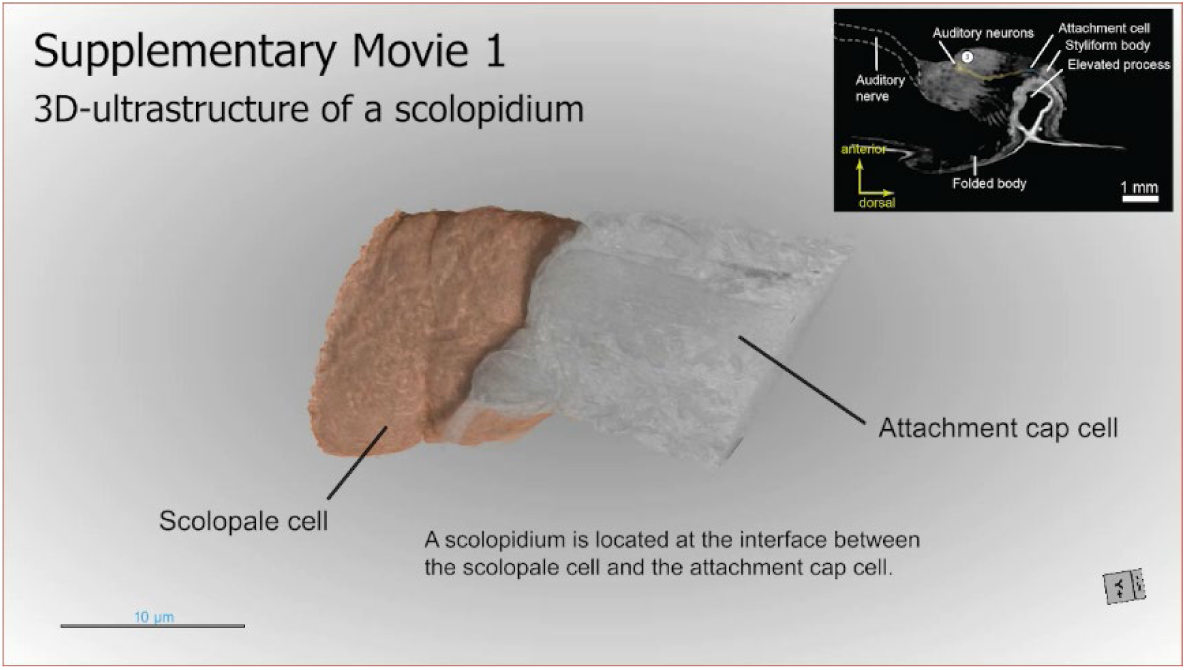
High-resolution 3D ultrastructure of a scolopidium using Focused Ion Beam-Scanning Electron Microscopy (FIB-SEM). Example of the 3D ultrastructure of a scolopidium with annotations^56^.

**SUPPLEMENTARY MOVIE 2.**
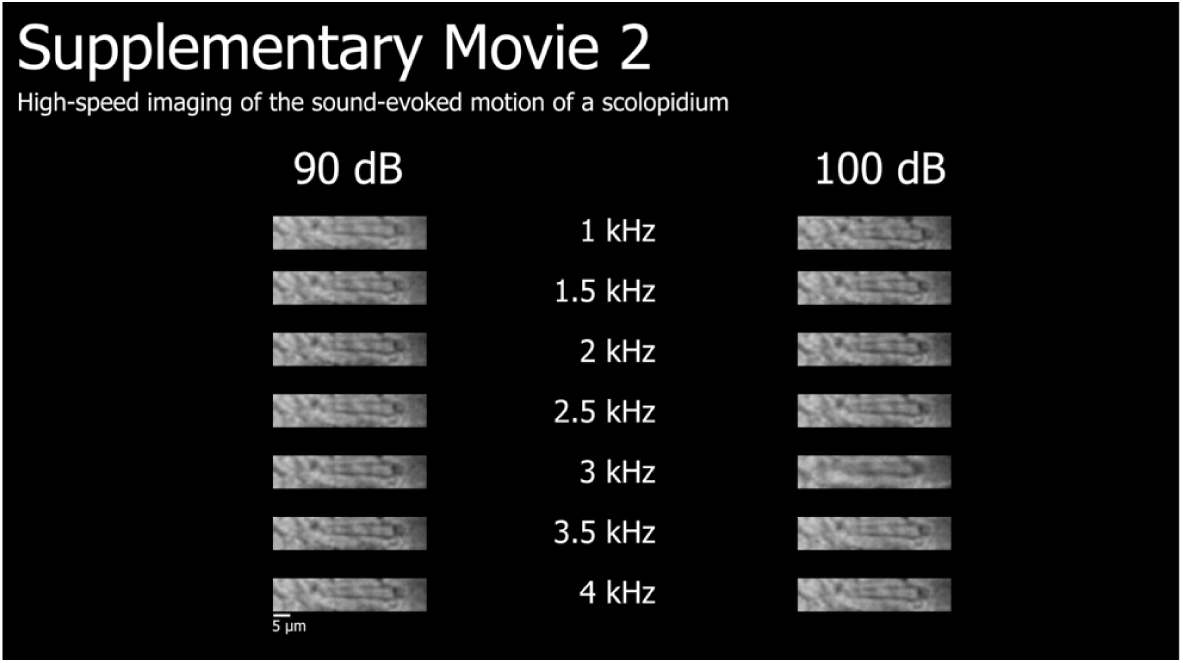
High-speed imaging of the sound-evoked motion of the scolopidium. Each scolopidium motion is played at a different frame rate. The frame rates can be found with additional movies and their metadata is in a public depository Zenodo^56^. This supplementary movie is based on the example Movie No. 1^55^.

**SUPPLEMENTARY MOVIE 3.**
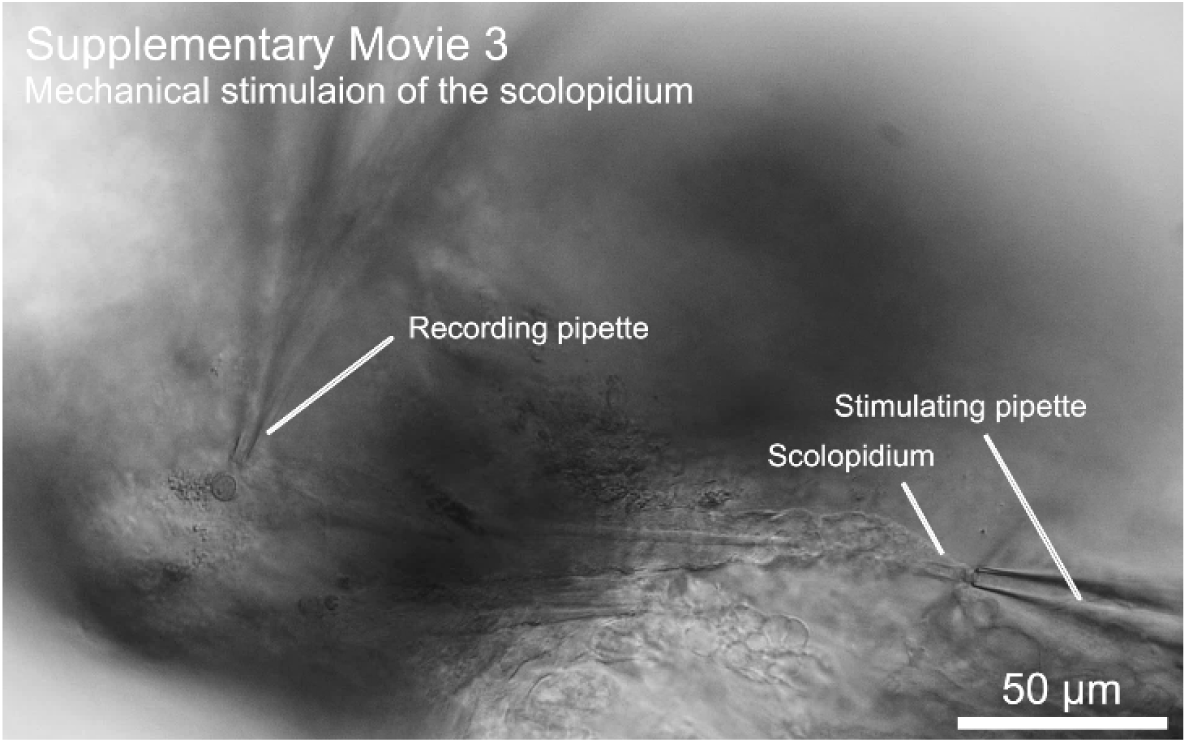
Mechanical stimulation of the scolopidium cap. The movie is played at 18.75 frames per second^56^.

